# Active teaching-learning methodology in a non-formal teaching environment as a tool to mitigate the ‘plant awareness disparity’ in public school students

**DOI:** 10.64898/2026.10.09.757080

**Authors:** Daniel Almeida Miron Soldi, Isabella Mesquita de Oliveira, Fernando Silvio Cavalcante Pimentel, Daniel Baron

## Abstract

Currently, a significant proportion of young people are discouraged by the teaching of botany and, consequently, show little interest in preserving local biodiversity. On the other hand, with the recurrent use of digital technologies (DTs), young people have shown significant interest in virtual devices. Our project’s tested the hypothesis whether plants awareness disparity in public school students was overcome through active teaching and learning. Thus, the objective was to investigate the use of active methodology in a non-formal teaching environment (e.g., sensory garden, SG) in order to stimulate learning in plant organography based on its evaluation using digital games. Our proposal was investigated at the Lagoa do Sino *campus* (Buri/SP, Brazil) in partnership with the Public School System of the Municipality of Campina do Monte Alegre/SP, Brazil. Furthermore, students were divided into two teams, which applied qualitative methodologies, such as that one of these (‘Team A’) had “face-to-face” contact with the SG, using the ‘Problem-Based Learning’ (PBL) methodology, digital game and survey. Team B, on the other hand, will only have contact with the game. After collecting the data, we verified statistically significant differences between the teams using the Wilcoxon test and based on the latent traits evaluated. Given this, the literature points out that discouraging students influences their learning and, consequently, their education. In addition, our considerations are based on the latent trait ‘knowledge of botany’ and, thus, we reject our null hypothesis and accept our alternative hypothesis. On the other hand, based on the latent trait of plant perception, we cannot affirm the acceptance or rejection of the hypotheses. It is worth noting that future studies should have a larger number of participants than this research in order to better evaluate what is intended.

**Societal impact Statement:** Brazil presents a paradox: it is an agricultural powerhouse with booming production and vast biodiversity, crucial to its economy. However, a significant “plant awareness disparity” exists. The public shows little interest in botany, and the media neglects its importance. This lack of knowledge means people fail to appreciate plants’ economic value in sectors like pharmaceuticals and cosmetics, or their environmental role. Consequently, there is little concern for preserving local biodiversity. Despite having the world’s richest flora, this motivational gap prevents society from valuing its vital botanical heritage.

## 1. INTRODUCTION

The school environment can act as a facilitator in promoting behaviors that will be reflected in their adult lives (Bezerra, 2017). These years (adolescence) are key periods of significant changes, both biological and social. The World Health Organization (WHO) has pointed out that the formation of ‘positive’ experiences and education in childhood/adolescence is fundamental to their development as citizens, in order to structure well-educated/informed individuals who can contribute satisfactorily to society (Langford et al., 2015). In Brazil, according to the ‘National Common Core Curriculum’ (BNCC), high school students have educational content related to plants, in which learning about the *Eukarya* domain (formerly the ‘Plant Kingdom’) and related subjects is included in the teaching axis focused on natural science subjects (Brazil, 2018).

The study of botany is the branch of science that studies plant diversity, as well as organographic definitions of plants and their variations (Perelman; Puhl, 2023). Botanical organography, also known as ‘external plant morphology’, studies the relationships between plants and their surrounding environment, their origins, as well as their changes and adaptations. This important area of botanical study investigates/describes aspects related to their forms and structures, which can be classified as ‘vegetative organs’, such as leaves, stems, and roots, or classified as ‘reproductive organs’, such as flowers, fruits, and seeds (Reis et al., 2016). It should be noted that Brazil has the greatest biodiversity in the world, which makes it unique in that it has biomes rich in different species of flora on the planet, such as the Amazon, Caatinga, Cerrado, Pampa, Atlantic Forest, and Pantanal biomes (Oliveira et al., 2012; Ulloa Ulloa, 2017; Santiago, 2018).

In the 21^st^ century, a considerable number of young students in public and/or private school systems show little interest in learning the content taught in botany classes (Da Silva; Feletti; De Oliveira, 2016). With the recurring use of digital technologies, society, especially adolescents, has become interested in visual digital devices (e.g., tablets and smartphones, generically described as “screens”) as opposed to interests dedicated to the natural environments that surround them, especially with regard to plant individuals, which are considered less “attractive” when compared, for example, animals (Fabeni, 2015; Salatino; Buckeridge, 2016). Thus, we must take into account the importance of directing botanical content to school-age youth.

In general, this national/international phenomenon is generically described as ‘botanical blindness’ or, as we will adopt in this project, ‘botanical imperception’ (Parsley, 2020; Ursi & Salatino, 2022). Given this, we can say that this phenomenon is a ‘latent trait’, that is, in order to assess whether individuals have higher perception ratings than others, a set of indices/parameters (such as a set of responses) is needed to analyze the desired measurements (Bichi; Talib, 2018). Among teachers and researchers focusing on botany, there is almost unanimous agreement that there is a notable ‘lack of interest’ in the ‘plant world’ among young students, that is, people do not devote sufficient attention (importance) to plant individuals (with very few exceptions), either in formal teaching environments (e.g., classrooms) or in their surroundings/daily lives (squares, parks, forests, among others) (Wandersee; Schussler 2002).

Unequivocally, a series of publications report that for people to respect and/or preserve and/or conserve and/or practice the proper use of natural resources, especially with regard to plants, it is essential to ‘know them’ (Salatino; Buckeridge, 2016; Neves; Bundchen; Lisboa, 2019; Dias *et al*., 2020). In addition, in order to stimulate learning in botany, the use of teaching tools that are attractive to young people and adolescents is becoming increasingly important. One promising and relevant strategy (often already established in other areas of knowledge) is the adoption of active teaching-learning methodologies (Roman, 2017).

Bearing in mind that learning occurs when students are motivated by a reason, which is often translated into a need, the use of new technologies (e.g., smartphones, apps, software, gamification) as methodological tools in the teaching and learning of topics related to botany can make the study of this important curricular component attractive and meaningful. In this line of reasoning, active teaching-learning methodologies fit perfectly as methods to overcome the “resistance” that the overwhelming number of HEIs (Higher Education Institutions) present in adopting teaching strategies demanded by the current moment in our society, for example, the problem of the lack of popularization and consolidation of sustainable techniques in the field, an “opportunity” for research and teaching. According to Bacich et al. (2015; 2017), research in the fields of education, neuroscience, and psychology proves that the learning process is unique and different for each human being, so that active methodologies encompass a conception of the teaching process that considers the effective participation of students in the construction of their learning, valuing the different ways in which they can be involved in this process so that they learn better, at their own pace, time, and style. Consequently, this method will awaken their autonomy and assist them in decision-making, whether individual or collective (Guarda, 2023). In addition to the intellectual gains mentioned above, the adoption of alternative teaching methods points to the possibility of innovating experiences, inside and outside the classroom, so that students of the ‘digital culture’ (21st-century students) feel stimulated to learn. When reflecting on the recent scenario, we refer especially to the 2020-22 triennium, when all of humanity faced the health crisis caused by the COVID-19 pandemic, which forced HEIs to seek solutions and/or didactic-methodological alternatives in the face of the compulsory closure of their physical teaching and research spaces. In the direction of this “new normal,” we observe a growing demand for educational expertise in the term “active methodology” (Irala; Blass; Ribeiro, 2020). What is understood as active methodology encompasses numerous possible teaching methodologies, including (but not limited to) “problem-based learning” (PBL) (Yew; Goh, 2016; Fernandes, 2021) and “design thinking” (DT) (Kenny et al., 2021; Lahiri; Cormican; Sampaio, 2021). Consequently, if any of these different teaching methods are applied, they will certainly stimulate student autonomy and assist them in decision-making, whether individual or collective (Guarda, 2023).

Digital technologies (DT) are part of social learning, so internet accessibility in schools is significant for motivating and innovative education (Pimentel, 2015). Educational institutions are deficient in this regard and considered incomplete because they do not prioritize technologies, which society is fully immersed in. That is, despite technological advances and easy access to information today, educational institutions have had difficulty keeping up with these developments and using them to their advantage. Face-to-face-digital-virtual teaching is a reality in the 21st century (Vetromille-Castro; Kieling, 2021).

The knowledge acquired through active methodology throughout the academic journey is relevant when it is aligned with the desires and objectives of each individual. Students who understand in advance that this information will help them, directly or indirectly, in their learning will certainly be encouraged to take ownership of the subject/content. Thus, it is important to consider the interests and needs of each student, something considered decisive for advancement in their education/academic life (Bacich, 2017). The ‘Problem-Based Learning’ (PBL) methodology essentially refers to the construction of knowledge through a problem or challenge that must be understood or solved, that is, its focus is on practical experiences. Furthermore, this strategy aims to incorporate different educational strategies to optimize student-centered learning outcomes so that students can acquire knowledge (Filipenko; Naslund, 2016).

In addition to contact between nature and humans, the use of digital technologies (DT) can, at first, distance young people from healthy visual/face-to-face contact with natural landscapes. However, there are ways to get around this problem, especially with the combination of DT and active teaching-learning. The rise of virtual technologies is driving the reinvention of educational environments and requires a new learning format. These effects generate opportunities for emerging teaching options, such as games in the educational context, or ‘digital games’. The use of games is becoming an important strategy in stimulating students to learn, i.e., digital games will become an important tool for sparking interest in botany based on something that is familiar to them and/or belongs to their ‘reality’. The use of digital games is based on the premise of proposing challenges, stages, problems, dealing with failures, and taking risks, so that these requirements “catch the attention” of students, providing them with a creative, imaginative, and curious experience (Cotta Orlandi et al., 2018). In parallel with the above, active teaching methods, such as the so-called ‘Bloom’s Taxonomy’ (Forehand, 2010), are presented as a classification system used to infer the level of cognitive performance of a given subject studied (West, 2023). The development of digital challenges can address teaching objectives based on their structure, such as ‘memorize,’ ‘understand,‘ ‘apply,’ ‘analyze,‘ ‘evaluate,’ and ‘create’ (Oliveira; Leite, 2024).

### 1.1 Societal impact statement: a case study in Brazil

We interpret the Brazilian scenario regarding botanical imperceptibility as paradoxical. On the one hand, Brazil is a country with an agricultural vocation given its favorable natural conditions for food production and, consequently, investments in the agricultural sector. For example, in Brazil, a significant part of agricultural production grew by 15.1% between 2022 and 2023, which boosted the national gross domestic product (GDP). During this same period, exports grew by 9.1%, an increase that stems from the main commodities produced in our country, such as corn (*Zea mays* L.) and soybeans [*Glycine max* (L.) Merr.] (Brazil, 2024). In this scenario, the country has competitive advantages in terms of the use of natural resources in food production, such as large areas of arable land, regions with different microclimates, and one of the largest freshwater reserves on the planet. On the other hand, there is a notable lack of interest among the population in understanding Brazil’s potential in matters of national interest, such as food security. This clear contrast is directly reflected in the neglect of the media and major news outlets (with very few honorable exceptions) in disseminating and popularizing topics that highlight the importance of the study of botany (Fachinello, 2024). In addition, the discovery of new natural molecules from Brazilian fauna and flora is of significant importance to the economy in different areas, such as pharmaceuticals, food, and cosmetics.

The scientific opportunities developed in the country are promising thanks to the biodiversity of Brazil’s genetic and natural heritage. For example, the state of São Paulo accounts for 32% of Brazil’s GDP and has a population of around 45 million, which accounts for 44% of the science developed in the country (Fapesp, 2017; Dos Santos; Silva; De Faria, 2023). The neglect of ‘knowledge’ about plants leads the population to not care, for example, about the preservation of local biodiversity and the importance of these living beings to society (both from an economic and environmental point of view). Although our country is considered to have the greatest biodiversity of flora on the planet, the lack of motivation to learn about plant biology contributes to plant awareness disparity.

Thus, we tested the hypothesis that the disparity in plant awareness among public school students can be overcome through active teaching and learning. Based on our “research question,” the details are as follows: (i) the disparity in plant awareness among public school students will not be mitigated through active teaching and learning [null hypothesis (“H0”)]. If “H0” is rejected, we will accept the alternative hypothesis (“H1”): the disparity in plant awareness among public school students will be mitigated through active teaching and learning. Thus, our objective is to investigate whether there is a correlation between knowledge of external plant morphology and botanical perception.

## 2. MATERIAL AND METHODS

### 2.1. Target audience

We evaluated 39 students from the public school system regularly enrolled in the 9^th^ grade of elementary school (aged between 13 and 15 years). In general, we assumed that the total number of students directly involved in the activities would represent a heterogeneous sample, even though the literature does not limit and/or recommend an exact number of students, given the countless scenarios found in public school classrooms. In addition, in partnership with the school administration (principal, vice principal, coordinator, scholarship candidate student, and teacher responsible for the proposal), we decided to divide the students into two teams, with 20 students in the control group and 19 in the treatment group.

#### 2.1.2. Formal teaching environment and informal teaching environment

The activities were carried out in two different locations. At the public school (formal teaching environment), active teaching activities such as “team-based problem solving” (PBL) and “digital games” were implemented. At the UFSCar Lagoa do Sino campus, students visited the sensory garden (SG) belonging to the ‘Experimental Garden and Forest Seedling Nursery’ of the Lagoa do Sino campus of the Federal University of São Carlos (UFSCar), located in the municipality of Buri/SP, a space called the sensory garden (non-formal learning environment) for the application of questionnaires. The SG has a total area of 54 m² (6 m x 9 m) and contains six small beds (SB), one central circular bed (CB), and one large bed (LB), all of which are 1.5 m high and constructed from reused materials (supplementary material, see Figures 1 and 2). We used 11 plant species (2 repetitions/individuals for each species) so that students could observe their different organographic characteristics (plant morphology). Below is a general compilation of the species supplementary material (Table 2).

#### 2.1.3 Team formation

The number of the students was 39 (thirty-nine), distributed across 4 (four) grade classes unrolled at High School, who were randomly selected into two teams, ‘A’ and ‘B’ (control team). The students had contact with SG during guided tours (activities carried out on site at SG) at ‘Team A’, besides participating in the active PBL methodology and contact with the game. On the other hand, students had contact plant organography classes in formal teaching environments according to the BNCC at their own school at Team B.

### 2.2. Experimental dynamics

#### 2.2.1. Application of the active teaching-learning methodology of the ‘PBL’ type (Problem-Based Learning)

We performed the teaching-learning methodology to ‘Team A’ students in the classroom according to Santos, (2021). Thus, the PBL theme was proposed in a ‘fictional case’ of forensic botany, regarding the structures and species that the students later encountered in SG. Our study challenge was to find the perpetrator of a ‘crime’ based on clues and evidence found during the investigation of the case. In this way, the students took on the role of “detectives”, encouraging them to assume leadership positions and engage in teamwork to solve the problem. The dynamics adopted were as follows steps (supplementary material).

### 2.3 Digital game application

We elaborated the game content according to ‘Bloom’s Taxonomy’ (Forehand, 2010) with minimal adjustments, which adopted the following classifications: (i) Perception of memorization; (ii) ‘Perception of understanding’, and (iii) Perception of the application of the content covered in order to facilitate the content of botany. Thus, the game content is related to the plant morphology, such as plant organs (leaf, stem, and root), plant reproductive structures (flower, inflorescence, fruit), botany family, genus, and species, as previously discussed with PBL and, subsequently, SG. Below, we present a complete description of each phase of the game, except for the evaluative approaches described in the section ‘Experimental evaluations’ (Supplementary material - Table 3).

### 2.4 Experimental evaluations

#### 2.4.1 Survey

In our study, we applied two ‘diagnostic questionnaires’ based on the pioneering study by Prestes *et al*. (2020) with modifications. The aim was to identify the desired Latent state-trait, such as “perception of plant species” and “knowledge of botany”. The ‘First and Second survey’ evaluated and compared the responses on the botanical content of Teams ‘A’ and ‘B’, based on the ‘Likert’ scale, which they classified with numerical scores on a scale of ‘1 to 5’ (1 = Strongly disagree, 5 = Strongly agree) (Adamsen; Rundle-Thiele; Whitty, 2013; Harpe, 2015). Therefore, we verified the progress of students who had contact with the active teaching methodology compared to the progress of students who did not have contact previously with the experimental methodologies on subjects directly regarding the plant morphology. Both surveys are available at the supplementary material.

#### 2.4.2 Statistical analyses

During the students’ participation in the game, we evaluated student performance by summing Likert averages, i.e., a nonparametric test, which we used to analyze data from the ‘Team A’ (treatment group) and ‘Team B’ (control group). In addition, we used ‘internal consistency’, which means (Menon *et al.,* 2025). Concerning these surveys, we intended to evaluate the content of the questionnaire about the latent state-traits labelled “perception of plant species” and “knowledge of botany”. Finally, the collected data were analyzed using the Wilcoxon test based on the teams’ averages. All analyses were performed in the R program using RStudio (4.5.1).

## 3 RESULTS

The results of the latent state-traits investigated in our investigation are shown in Figure 3. Thus, we observed that for the ‘knowledge of botany’, the results for the students at ‘Team A’ (treated group) were consistent, based on the responses obtained from the students (*w* = 52, *p* < 0.0001). Thus, according to Cronbach’s alpha data (Menon *et al*., 2025) these results confirm that the students who previously participated in the theoretical basis and practical activities in botany outperformed the other students in ‘Team B’ (control group).

**Figure 3.**
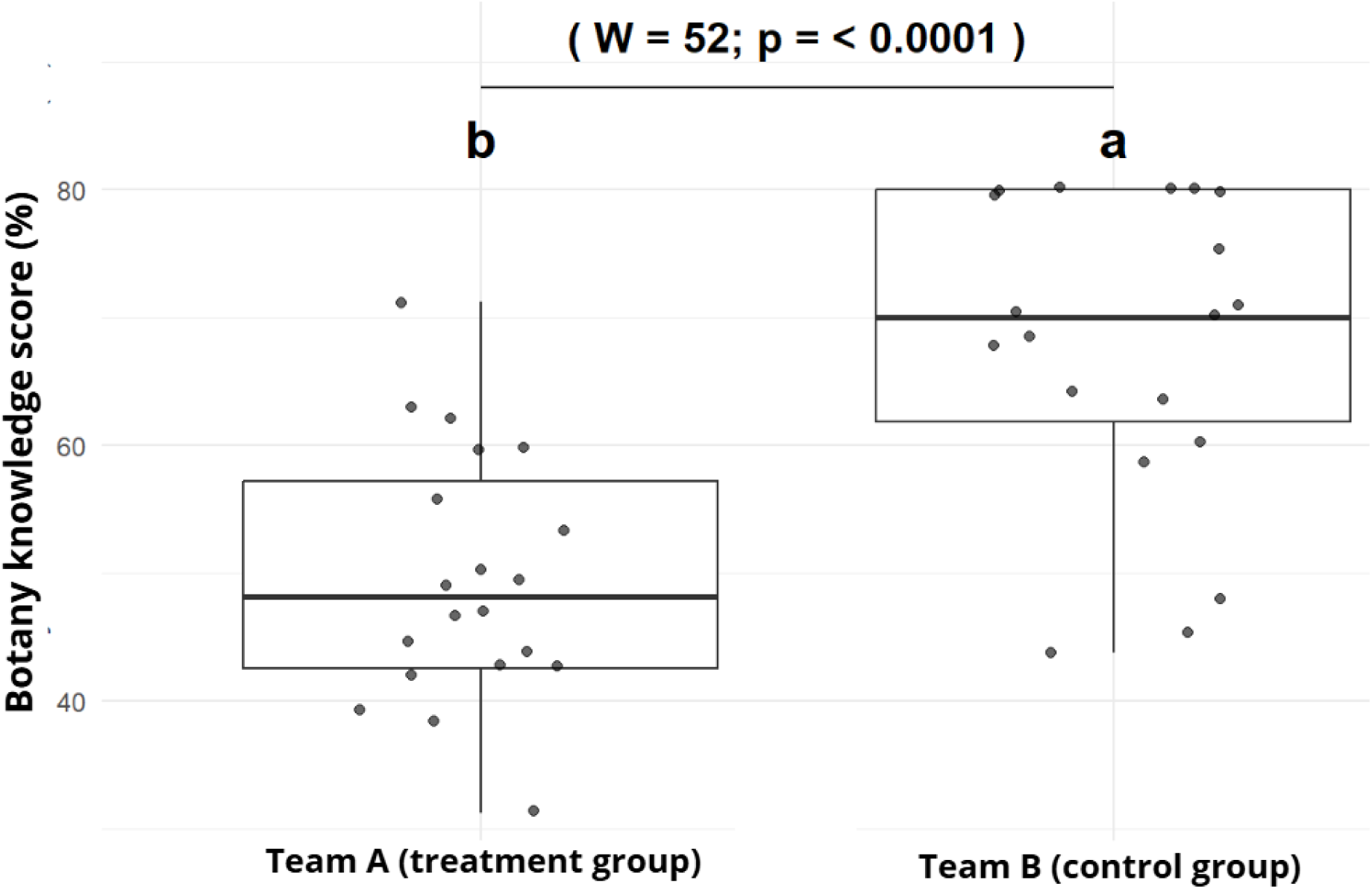
Median values for teams A and B in relation to the latent trait “botany knowledge score” showed a significant difference *p* = < 0.05 and a Cronbach’s alpha value of 0.789 (a satisfactory value according to the Cronbach’s scale).

As for the latent trait “perception of plant species” the results indicate that there is no internal consistency between the data, but there were statistical differences between the teams evaluated (Figure 4). The internal consistency in this second latent trait reveals that the reported statistical difference is not due to the questionnaire answered by the students, but rather to a myriad of other causes and/or situations.

**Figure 4.**
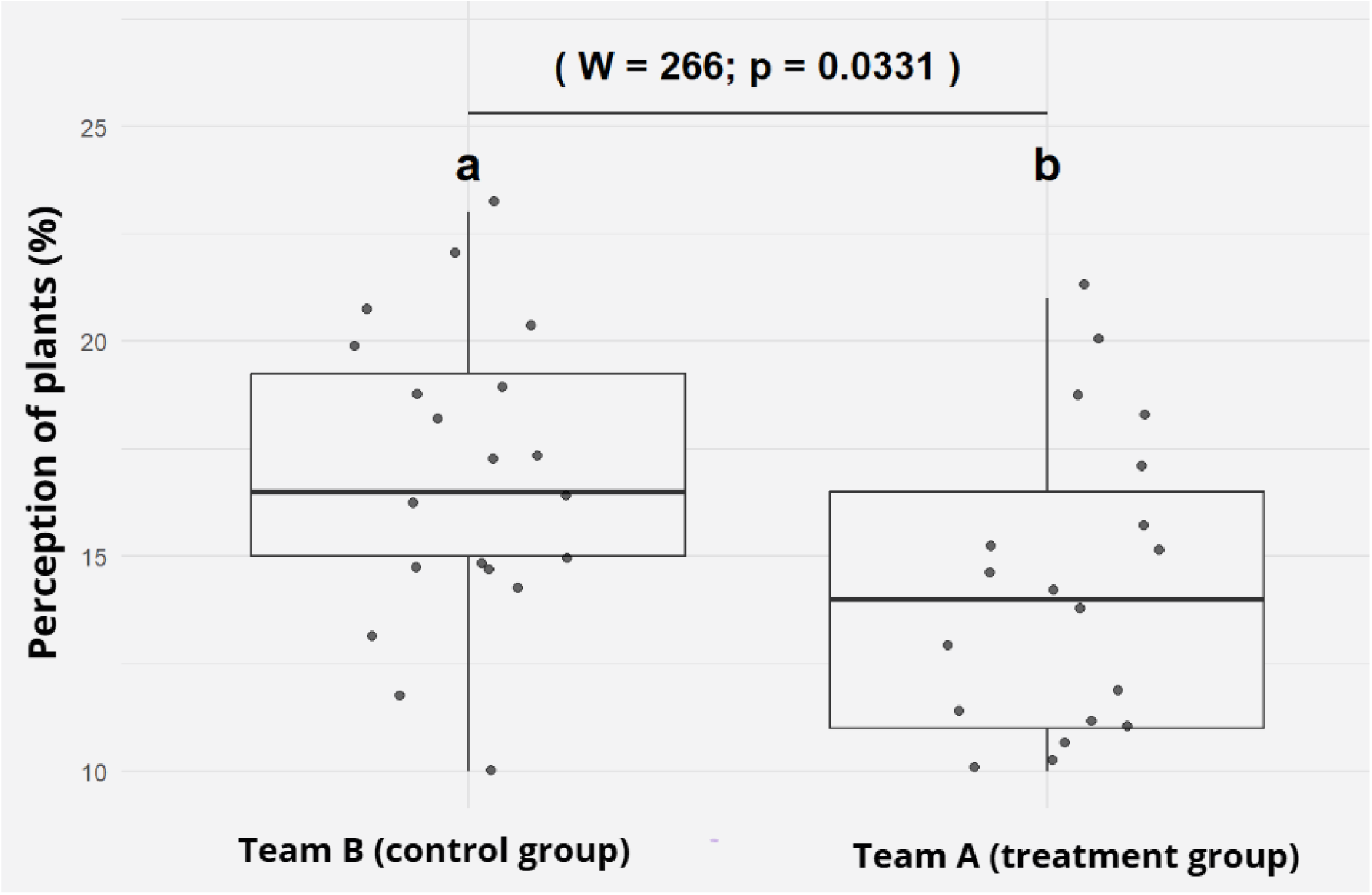
Median values for teams A and B in relation to the latent trait “perception of plants” which, using the Wilcoxon test, showed a significant difference *p* = < 0.05 and a Cronbach’s alpha value of 0.285 (unsatisfactory value according to the Cronbach scale).

Finally, when evaluating the Pearson correlation between the two latent traits evaluated (Figure 5), our results report that Team B (control group) maintained low values of intellectual gains in ‘botanical knowledge’, which is confirmed by their significant ‘botanical imperceptibility’. Team A (the treatment group), on the other hand, showed intellectual gains in their learning under “botany knowledge,” which consequently resulted in a significant decrease in “perception of plant species”.

**Figure 5.**
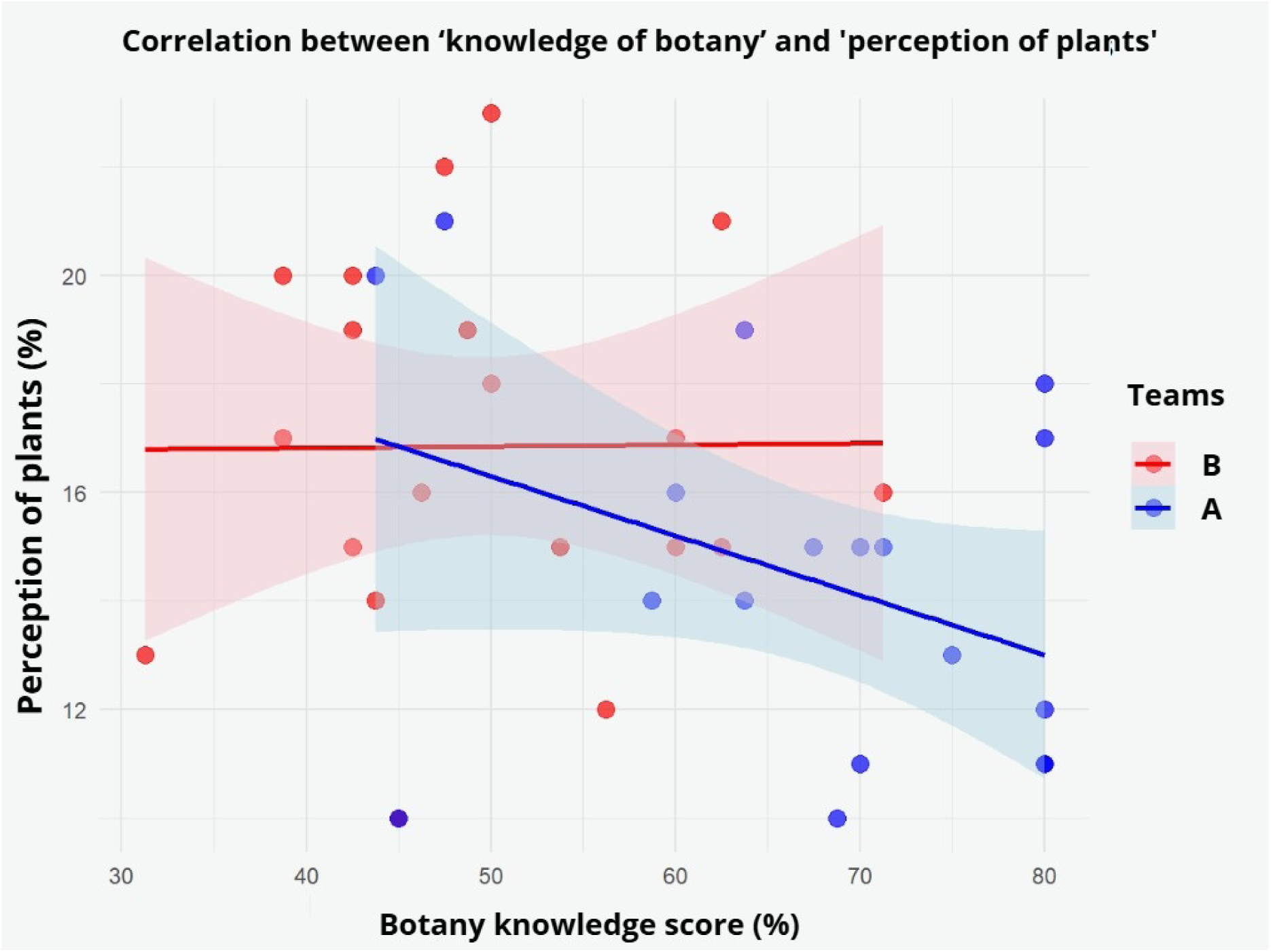
Pearson correlation between the two latent traits that showed statistical difference, in blue ‘team A’ and red ‘team B’, where the ‘x’ axis represents the latent trait of the botany knowledge score and the ‘y’ axis shows the levels of plant perceptions.

## 4. DISCUSSION

In general, the adoption of active teaching and learning proved to be effective in overcoming plant awareness disparity among young students in the public school system. This finding is due to the fact that, through the diagnosis of the latent trait that we arbitrarily defined as ‘botany knowledge,’ it confirms the learning of the didactic content addressed on ‘external plant morphology’ (plant organography). Thus, we show that initial limitations in learning botany can be reliably overcome through active teaching-learning strategies, which do not belong to the daily and/or periodic routine of the students evaluated.

In general, the use of active teaching-learning strategies is not new and is significant in Brazilian contexts. For example, Borochovicius and Tasson (2021) report that the use of problem-based learning (PBL) optimizes learning for elementary school students. In this way, students become protagonists of their own learning, reflecting on, synthesizing, and directly/indirectly applying the content covered, unlike conventional classes in formal teaching environments, where attention is focused exclusively on the teacher and there is no “exchange” between student and teacher. Authors such as Delgado (2022) and Oliveira (2024) affirm that academic gains are due to the motivation provided by the topic/subject being discussed, as well as the stimulus to solve a challenge or problem. This finding about the stimulus needed to motivate students to learn corroborates what we use as an “interest trigger” activity through the use of a fictional forensic criminalistics case, based on the survey of botanical characteristics obtained at the “crime scene” to solve the case. In addition, authors such as De Almeida, David, and Ribeiro (2022) affirm that items belonging to the student’s daily life (e.g., the internet and online games) are of immediate interest and application in their daily lives.

According to recently published literature reviews, such as Costa (2023) and Brasil (2021), the use of teaching strategies has a direct impact on students’ neurological state. These same authors report that, when investigating brain activity in students who had direct contact with active methodologies, they showed a significant difference in intellectual activity when compared to students who did not have the same direct contact with active teaching (Brasil, 2021). In addition, Grossi, Lyra, and Silva (2024) state that active teaching-learning techniques resulted in improvements in studying. Yoo (2012) and Amaral; Guerra (2022) state that learning in the human brain presents attributes in the formation of connections between nerve cells that occur through synapses and circuits of neurons connected and distributed in the central nervous system. In neurology, learning is interpreted through the existence of structuring and consolidation of connections in the space that separates cells, therefore, the effect of experienced sensations causes chemical and even structural changes in the brain.

On the other hand, students without prior contact with active methodologies rely exclusively on the “botanical foundation” derived from their educational interactions with public school teachers during their elementary school years. According to Ursi (2018), a significant portion of teachers did not acquire the solid and robust learning necessary in plant biology during their teacher training and/or, when they did, it was only in general biology, which inevitably compromises the transmission of information about botany to their students. Furthermore, general botany subjects, often marginalized and/or omitted from the curriculum, further aggravate the situation, given that these subjects are not part of their main area of training. Nevertheless, the extremely rare financial rewards (without real perception of remuneration), as well as unattractive and demotivating career plans, further reinforce the fact that teachers do not seek to improve and update their technical knowledge on the subject, which tends to aggravate the situation of demotivation for training and interest among future and potential teachers in the area of elementary education (Gatti, 2012; Brasileiro; De França; Da Silva, 2025).

In addition to this worrying scenario, it is worth noting that botanical neglect is present in the daily life of our current society. Authors such as Salatino and Buckrebridge (2016) conclude that it is important for plants to play a prominent role in the media and communications, especially at a time when the overwhelming majority of young people are connected to the global computer network and, thus, increasingly in contact with social media in our daily lives. Our study revealed that young people, when minimally stimulated, paid attention to organographic characteristics and showed interest in learning the basics of plant morphology in order to complete the proposed activities. Thus, our findings corroborate what some authors have previously stated, such as Bacich (2017) and Crestani (2023), that low interest in botany occurs until students find meaning in their learning.

As for the latent trait “perception of plant species,” it should be noted that, as we did arbitrarily for the previous latent trait, we listed a series of questions that we considered directly related to the topic. Our findings allow us to reflect that there is no way to assume what the real contribution of active methods was in overcoming this latent trait or, at most, allow us to speculate about it. For example, the implicit content addressed in our questionnaires was compiled arbitrarily by the authors, that is, there is no clear and precise indication in the literature of which question(s) are specific to address in a descriptive study on the difficulty of learning botany. Thus, we chose to apply questions based on the “practical experience of students’ difficulties in their daily lives” regarding the recognition and applications of plants in their daily lives. As for the number of questions used in each questionnaire, we adopted a number close to that used by Prestes *et al*. (2020), so that the quantitative assessment would not discourage respondents, and we did not emphasize questions with a higher degree of complexity for the same reason.

In general terms, the literature recommends that for each latent trait evaluated, a significant number of participants (between 20 and 90 respondents) should be investigated. It is worth noting that our research met this range of participants (39 respondents), although it was below our desired limit. Thus, the reality we investigated reliably reflects the overwhelming majority of public schools in Brazil, where student participation in ‘extracurricular’ activities is a tremendous challenge. In addition, other unique situations proved challenging in our school context, such as the physical space and electronic equipment (computers) for teaching activities, which limited the final number of student participants.

Furthermore, it is worth noting that the environment surrounding young students often does not favor their learning. According to Patto (2015), social issues such as the level of education of students’ parents, where students live, and economic class (middle and/or low) influence student learning, since these families do not have a positive structure in terms of preparing students to deal with the demands required/requested by educational environments. Other authors such as Souza; Nobrega; Freitas, (2019); Souza, (2025) report that there may be conflicts between the school environment and the family environment, given that families have a singular way of thinking and project their expectation that the family environment corresponds in the same way to the demands of schools. However, educational spaces become a training environment that should emphasize the individuality of students, but their actions end up reflecting a homogenized way of treating students and their families.

Another bottleneck faced in research with public school students is the vast gap between higher education institutions and elementary schools. In Brazil, historically, these two “audiences” have had difficulty “communicating with each other” and, unfortunately, this observation results today in a reality in which research activities are rarely undertaken by elementary schools that seek interaction between these audiences. It should also be noted that a significant number of university study projects have never been implemented and/or completed “to satisfaction” in public schools (unpublished data) for a variety of reasons, whether due to cuts in public funding for education (Chaves, 2020), or “in and of itself” in the execution of projects, or due to the political/cultural polarization (Salata, 2018) underway in Brazil and around the world, so that all these factors, whether combined or not, will also deepen this “gap” that separates the different institutions.

## 5 CONCLUSION

Given the above, based on the latent trait ‘knowledge of botany’, we reject our null hypothesis and accept our alternative hypothesis that active teaching-learning mitigated botanical imperceptibility in public school students. However, with regard to the other latent trait, ‘perception of plant species’, although the teams showed a statistical difference between them, the Cronbach’s alpha value did not show internal consistency between the questionnaires, so we cannot accept or reject our respective H0 and H1. As future perspectives, we suggest that although there is a difficulty in the number of participants (students), there is a considerable number for conducting the study. Of course, there are variables that are not within the authors’ control, such as the support of those responsible for the students’ participation, the school environment, and social media that encourage these students to contribute to topics related to plant morphology.

## Supplementary material - 1

**Figure 1.**
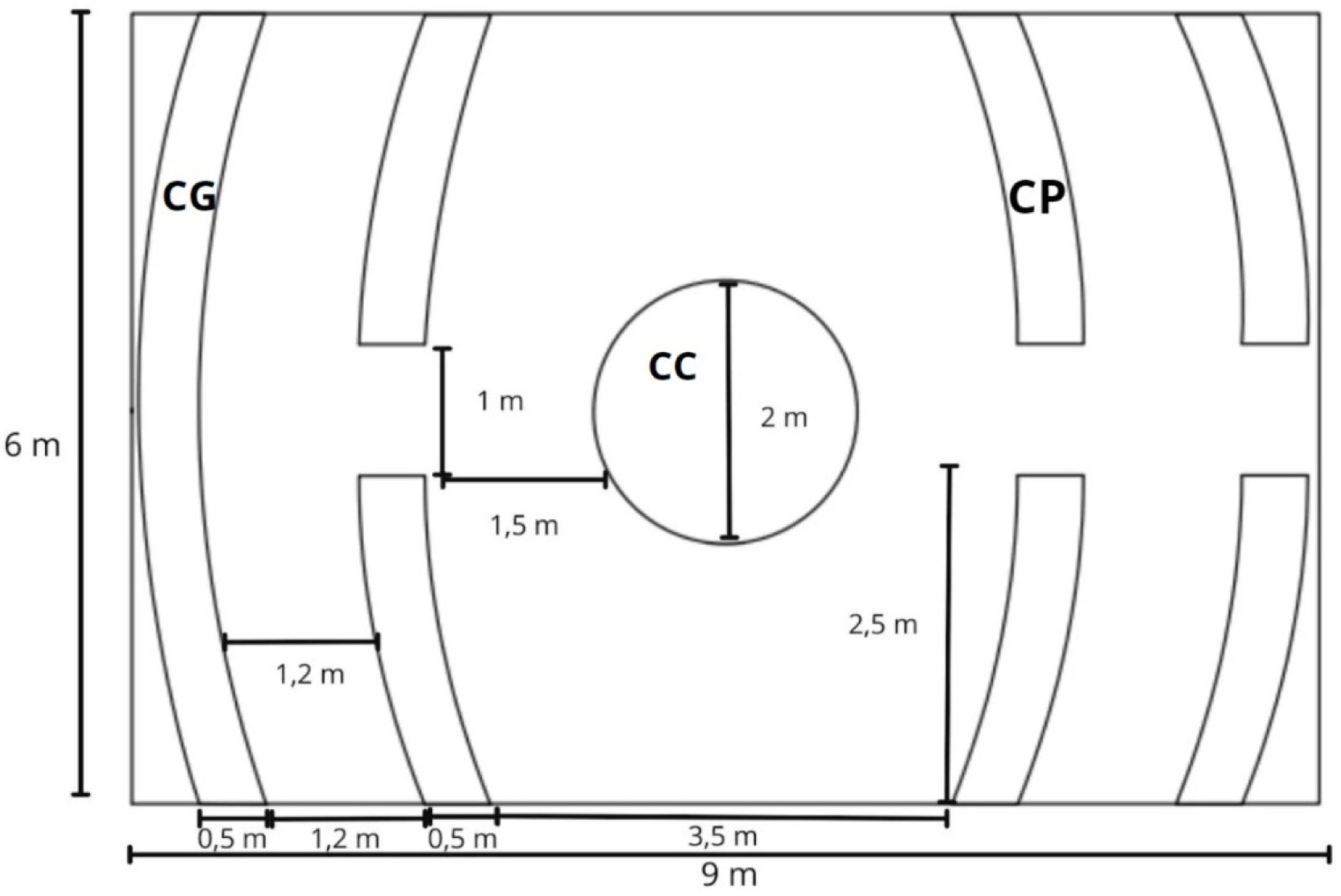
- Dimensions of the floor plan/planimetric/altimetric/two-dimensional sketch. Measurement of the informal teaching space described as a Sensory Garden.

- Measurements of the small flower bed (**SB**):
  - Perimeter: 6 m
  - Clear height of the chip: 0.5 m
  - Size of the chip to remain buried: 0.3 m
  - Area: 1.25 m²
  - Volume: 0.7 m³
- Measurements of the central flowerbed (CB):
  - Perimeter: 6 m
  - Radius: 1 m
  - Clear height of the chip: 0.5 m
  - Size of the chip to remain buried: 0.3 m
  - Area: 3m^2^
  - Volume: 0.6m^3^
- Large flowerbed (LB) measurements:
  - Perimeter: 7m
  - Clear height of the chip: 0.5 m
  - Size of the chip to remain buried: 0.3 m
  - Area: 3m^2^
  - Volume: 1.5m^3^

## Supplementary material - 2

**Table 1.** List of permanent materials and consumables to be used for the construction of the SG.

| Permanent Materials | Quantity | Unit/measurements |
| --- | --- | --- |
| Galvanized steel wire | 1 | 40 meters |
| Tape measure | 1 | 10 meters |
| Extension cord/ruler | 1 | 10 meters |
| Aquarium pump | 1 | unit |
| Waterfall fountain | 1 | unit |
| Hoe | 3 | units |
| Eucalyptus wood chips | 300 | units |
| Trowels | 50 | units |
| Mineral fertilizer (N, P, K) | 10 | kg |
| Fine sand | 30 | kg |
| Gravel | 50 | kg |
| Organic substrate<br>(compost) | 50 | kg |
| Plants used (Table 2) | 22 | units |

## Supplementary material – 3

**Figure 2.**
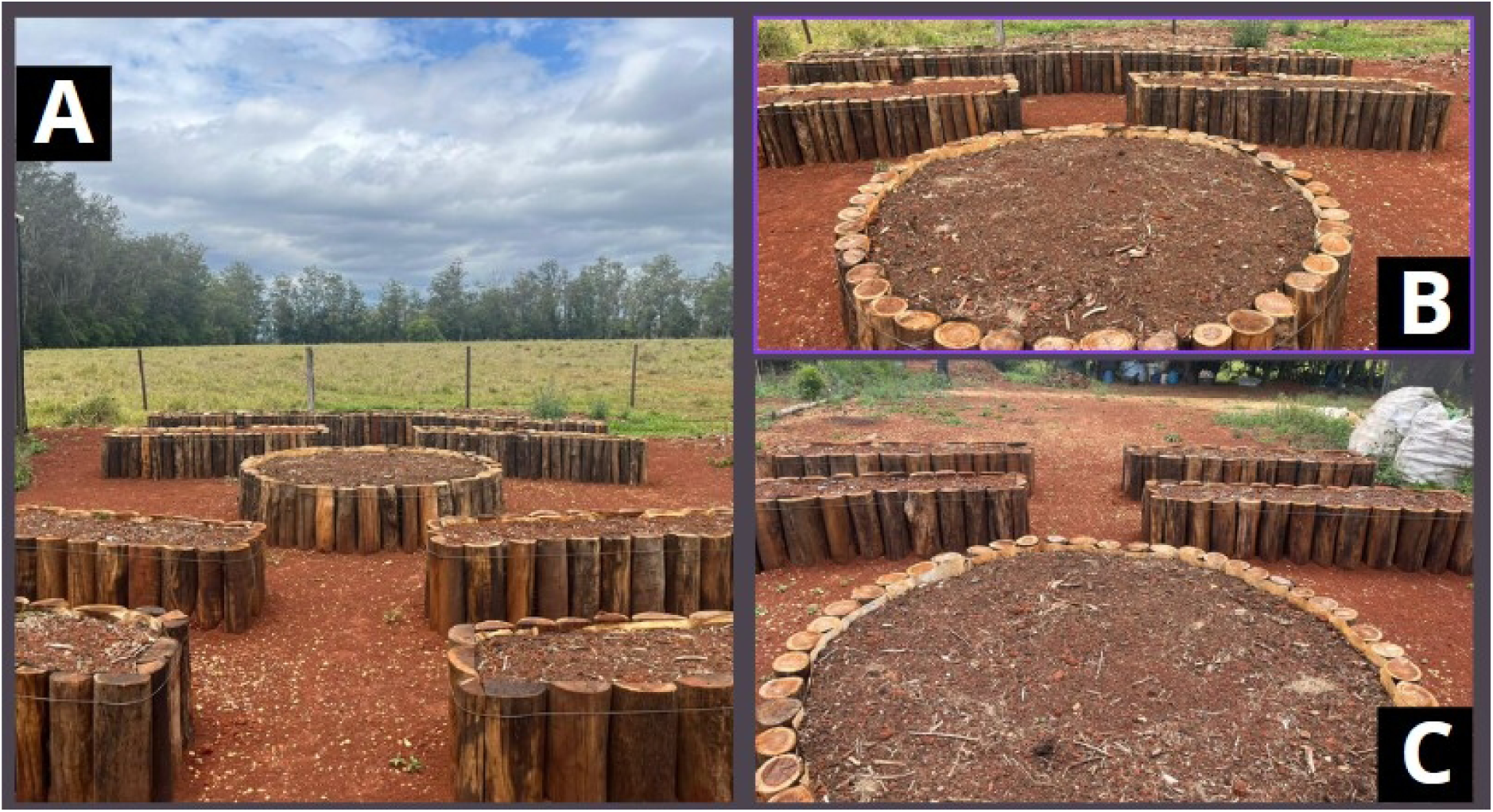
Overview of the construction and installation of the planting beds that make up the sensory garden (non-formal teaching environment) located in the ‘Experimental Garden and Forest Seedling Nursery’, belonging to the Federal University of São Carlos (UFSCar) Lagoa do Sino campus (Buri/SP) with geographical coordinates 48°31’42’‘W 23°36’05’’S. (**A**) Frontal view of the SG and its beds, SB, CB, LB; (**B**) View from the CB; (**C**) View from the center of the SG, overlooking the four SB, after CB. **Image credits: D**. A. M. Soldi

The beds are filled with a mixture of soil classified as typical Eutrophic Red Latosol with a clayey texture (53%) and compost (vermicompost).

## Supplementary material – 4

**Table 2.** Plant species to be used in the installation of the sensory garden located in the ‘Experimental Garden and Forest Seedling Nursery’ belonging to the Federal University of São Carlos (UFSCar) Lagoa do Sino campus (Buri/SP).

| Common name | Scientific name | Botanical family | Sensory attributes | Authors |
| --- | --- | --- | --- | --- |
| rosemary | *Rosmarinus officinalis* L. | Lamiaceae | Smell | (Cordeiro et al., 2020) |
| azalea | *Rhododendron simsii* Planch. | Ericaceae | Sight/Hearing | (Borges; Paiva, 2009) |
| boldo | *Plectranthus sp.* | Lamiaceae | Touch | (Campelo; Santos; Souza, 2021) |
| lemongrass | *Cymbopogon citratus* (DC.) | Poaceae | Touch | (Borges; Paiva, 2009) |
| nasturtium | *Tropaeolum majus* L. | Tropaeolaceae | Taste | (Machado; Barros, 2020) |
| chives | *Allium fistulosum* L. | Amaryllidaceae | Taste | (Borges; Paiva, 2009) |
| christmas cactus | *Schlumbergera truncata* (Haworth) | Cactaceae | Sight | (Borges; Paiva, 2009) |
| guaimbê | *Philodendron selloum* Schott. | Araceae | Sight/Hearing | (Cordeiro et al., 2020) |
| mint | *Mentha piperita* L. | Lamiaceae | Taste/Smell | (Machado; Barros, 2020) |
| basil | *Ocimum basilicum* L. | Lamiaceae | Taste/Smell | (Cordeiro et al., 2020) |
| oregano | *Origanum vulgare* L. | Lamiaceae | Smell/Taste | (Machado; Barros, 2020) |
| bear's paw | *Cotyledon tomentosa* Harv. | Crassulaceae | Touch/Sight | (Machado; Barros, 2020) |
| parsley | *Petroselinum crispum* (Mill.) Nym | Apiaceae | Smell/Taste | (Machado; Barros, 2020) |
| thyme | *Thymus vulgaris* L. | Lamiaceae | Taste/Smell | (Machado; Barros, 2020) |

## Supplementary material - 5

A) Botanical imperceptibility is the lack of awareness of the importance of plants to society.

B) The function of the root is to anchor the plant to the soil and absorb water and nutrients.

C) In most species, photosynthesis is not performed by the roots.

D) The taproot has the appearance of a single, main axis.

E) Tree species have a “trunk” type stem.

F) A palm tree has a stipe type stem.

G) The stem is not the plant organ responsible for attracting pollinators.

H) The stem is the plant organ that connects the roots to the leaves, in addition to supporting the plant.

I) The leaf is not the organ responsible for protecting and dispersing seeds.

J) The leaf is the main organ responsible for photosynthesis.

K) Photosynthesis is the conversion of light energy into chemical energy.

L) Through photosynthesis, plants produce their own “food.”

M) In addition to their reproductive function, flowers are also responsible for attracting pollinators.

N) Flowers are not the organs responsible for supporting the plant.

O) Fruits are not responsible for anchoring the plant to the ground.

P) The fruit is the organ responsible for protecting and dispersing seeds.

## Supplementary material - 6

A) Are there currently any plants on the sidewalk and/or in front of your house and/or home?

B) Are the foods you consume in your daily meals (breakfast, lunch, and dinner) of plant origin (directly and/or indirectly)?

C) Do you usually observe the plants that are part of your daily commute to school (squares, gardens, parks, sidewalks, etc.)?

D) I consider it important to understand the role of plants in sustaining life on the planet.

E) I feel motivated to learn botany on my own.

F) I believe it is essential to know the importance of plants to society.

G) I consider the study of botany essential for the preservation of native species.

H) I believe that botany classes could be more dynamic with practical activities.

I) The study of plants contributes to the appreciation of natural resources.

J) Botany helps me to better understand health and well-being issues (food, medicinal plants).

## Supplementary material – 7

**Table 3.**
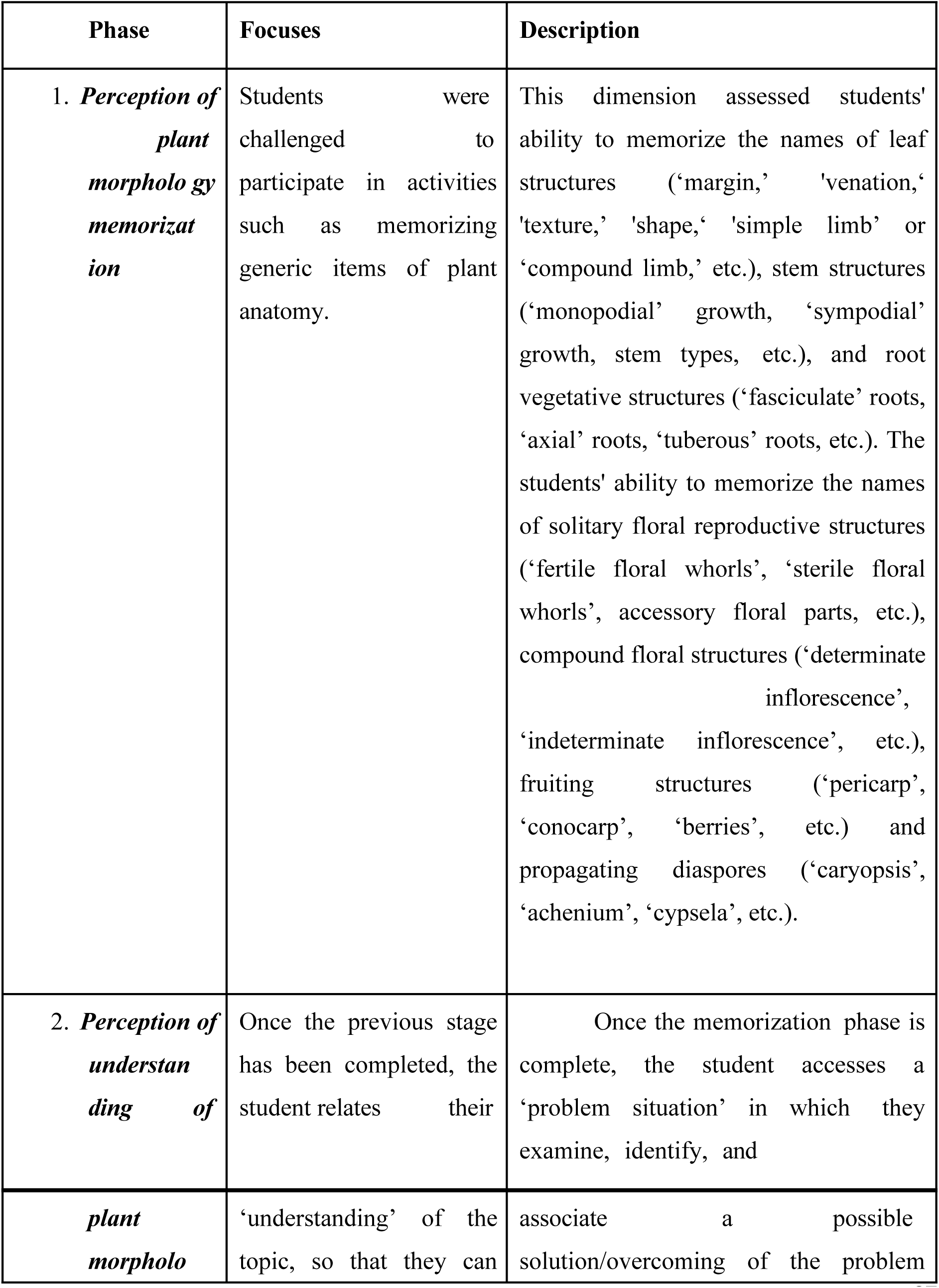
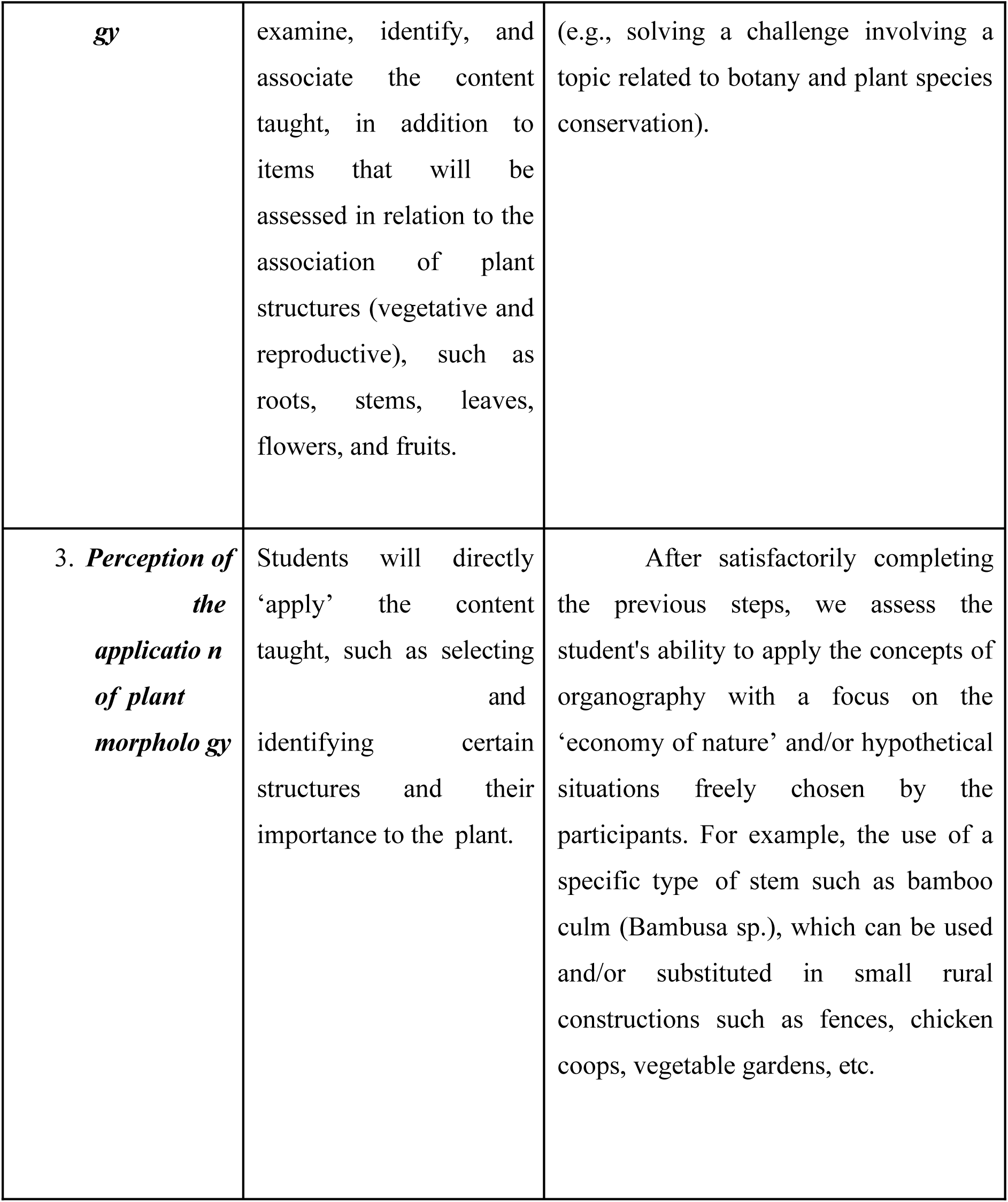
Overview of different phases applied to High School students according to ‘Bloom’s Taxonomy’ (Forehand, 2010) with minimal adjustments.

## Supplementary material - 8

### 2.2 Methodological approach

#### 2.2.1 Mandatory bureaucracy

Initially, public school students received general instructions on the entire experimental dynamic, as well as a detailed approach to the practical activities of the project, such as the students’ visit to the SG, application of the active methodology, and the online game focused on ‘plant organography’ content. In addition to these tools, we incorporated qualitative methods with ‘longitudinal approaches’. For example, we collected data on the participants ‘relationship with plants’, i.e., whether the students have frequent contact with external plant morphology (organography) outside the classroom or only in the classroom. The Brazil Platform serves as the unified national basis for the registration and monitoring of research projects involving human beings in Brazil, integrating the Research Ethics Committees System (CEP)/National Research Ethics Commission (Conep). The digital platform facilitates the submission, review, and monitoring of projects from conception to final approval, making the process safe and reliable for researchers, institutions, and research participants. In view of this, participants were made aware of the bureaucratic requirements of the Research Ethics Committee (‘CEP’) [CAAE: 89130825.7.0000.5504]. The students were duly informed that their participation is not mandatory and, if anyone feels uncomfortable and/or embarrassed to participate in the activity, they may, at any time and without prejudice, withdraw from participating in future activities. Furthermore, plant species belonging to Brazil’s natural genetic heritage and traditional knowledge (SISGEN) were digitally reported in the electronic system (https://sisgen.gov.br). The digital platform is available so that we have already reported the scientific name of the plant species used and their origin for registration.

## Supplementary material - 9

### Application of the active teaching-learning methodology of the ‘PBL’ type (Problem-Based Learning

Step 1. We presented the botanical investigation theme (“*unraveling the mystery and four suspects incriminated” for being the “possible perpetrators of the theft of a rare jewel”*). Students had to find evidence from plant morphology (*previously presented structures*) located on the suspects’ clothing;

Step 2. Based on the progress of the investigation, each of the sub-teams (between four and five students) was guided by marking a map and the plant species found in the region of the murder, as well as those that are not located near the crime scene. In addition, we encouraged the investigation and answered questions about the plant morphology, without compromising the solve of the problem, i.e., they did not address the issue of who was responsible for the murder, but only the botanical characteristics;

Step 3. In this last step, a debate was held on the case so that students could talk and discuss the types of stems, roots, leaves, flowers, and fruits found and relate them to the perpetrator suspects. The question of who committed the theft was asked based on the decisions made by the students who analyzed the entire dynamics, which included the plant morphology found on the clothing, as well as whether they were regarding the plant individuals found at the crime scene. Then, a definitive conclusion was found as to which of the four crime perpetrator suspects. Thus, the problem was solved based on plant morphology.

## Supplementary material - 10

**Figure 6.**
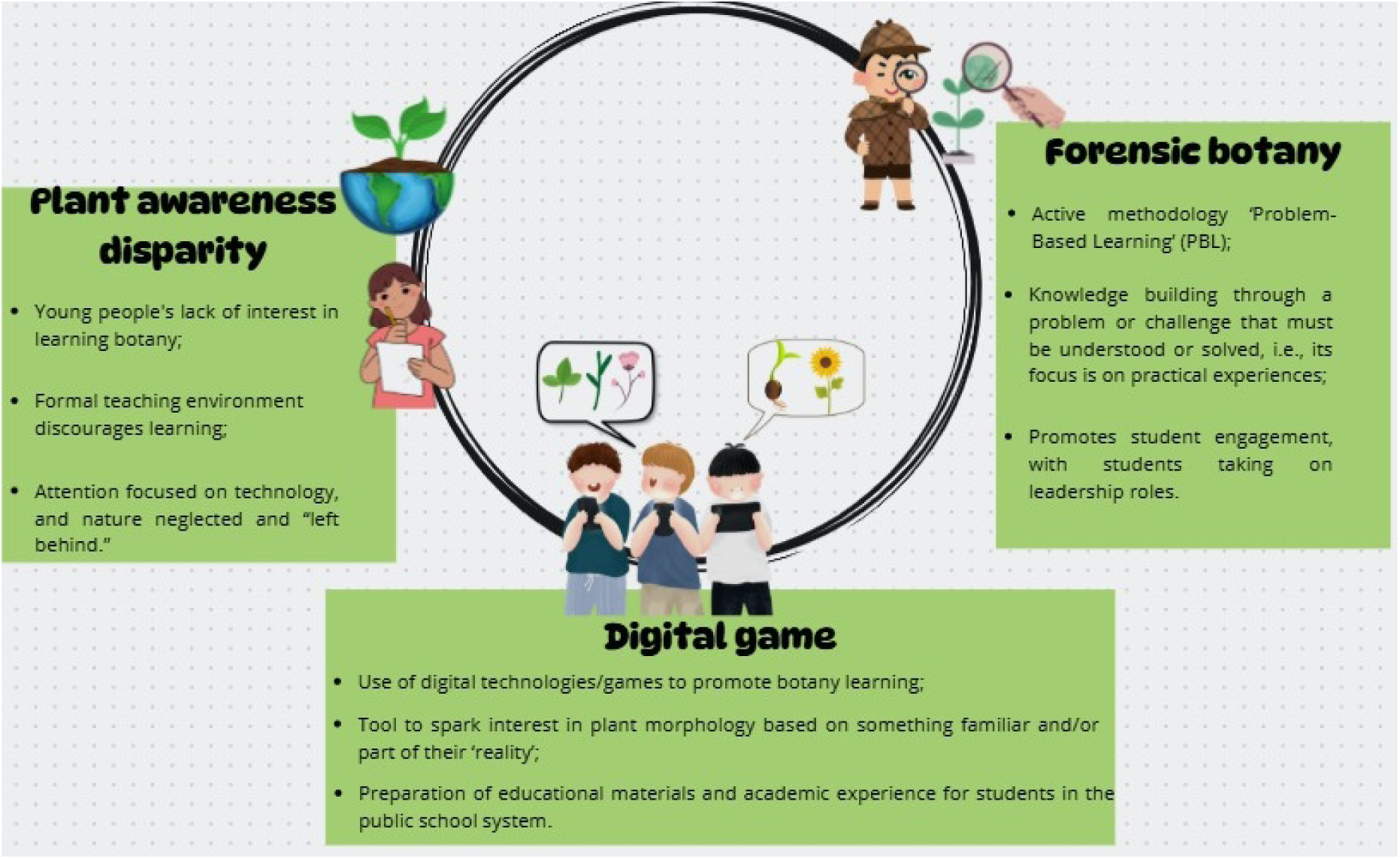
Mind map with the fundamental premises that justify the use of active teaching-learning methodologies in non-formal teaching environments. Image credit: D. A. M. Soldi.

